# High-Resolution, Long-Term Trends in Relative Abundance from Spatial Modeling of Continent-Wide Bird Counts

**DOI:** 10.1101/466425

**Authors:** Timothy D. Meehan, Nicole L. Michel, Håvard Rue

## Abstract

Continent-wide bird counts by community volunteers provide valuable information about the conservation needs of many bird species. The statistical modeling techniques commonly used to analyze these counts provide robust long-term trend estimates from heterogeneous community science data at regional, national, and continental scales. Here we present a novel modeling framework that increases the spatial resolution of trend estimates, and reduces the computational burden of trend estimation, each by an order of magnitude. We demonstrate the approach with data for the American Robin (*Turdus migratorius*) from Audubon Christmas Bird Counts conducted between 1966 and 2017, and show that aggregate regional trend estimates from the proposed method align well with those from the current standard method. Thus, it appears that the proposed technique can provide reasonable large-scale trend estimates for users concerned with general patterns, while also providing higher resolution estimates for others examining correlates of abundance trends at finer spatial scales.

## Introduction

Volunteers with the Audubon Christmas Bird Count (CBC) have been counting wintering birds across North America every year for the last 118 years (Dunn et al. 2005, Soykan et al. 2016). Population trends derived from CBC data, along with those derived from other large-scale monitoring programs like the North American Breeding Bird Survey (BBS, Robbins et al. 1989, Sauer et al. 2017), provide valuable information for understanding the conservation needs of North American bird species (Dickinson et al. 2010, Hochachka et al. 2012, Rosenberg et al. 2016).

The current, standard approach for generating trends from CBC data (Link et al. 2006, Soykan et al. 2016) was derived from methods originally developed for BBS data (Link and Sauer 2002, Sauer and Link 2011). The general approach is to assign counts in Canada and the US to one of up to 169 polygons or spatial strata, which are intersections of US states, Canadian provinces, and Bird Conservation Regions (BCR, Sauer et al. 2003). Then, treating each stratum as independent, a non-linear function is used to correct for the effect of observer effort on counts, and model the residual as a function of count circle, stratum, and year (Link et al. 2006, Soykan et al. 2016). These parameter estimates are used to derive a relative abundance index per stratum and year, and those indices are used to compute annual percent change per stratum across defined time periods (Link and Sauer 2002, Sauer and Link 2011).

The traditional CBC analysis provides robust long-term trend estimates from heterogeneous community science data across large spatial scales. By pooling count circles per stratum, this approach deals with the issue of count locations (here, CBC circles) haphazardly becoming active or inactive over the time series (Sauer and Link 2011, Soykan et al. 2016). Additionally, pooling produces a sufficiently large sample of counts to generate a reasonably robust count-effort correction function (Link and Sauer 1999), which is critical given the wide variation in count effort among count circles (Bock and Root 1981, Dunn et al. 2005). This approach produces a relative abundance index per year and stratum, which can be used to explore variation around long-term, log-linear trends, and can be summed across larger hierarchically nested strata, such as states, provinces, or BCRs, and used to calculate change in relative abundance at larger spatial scales. Producing annual abundance indices also permits summarizing abundance change between any desired pair of time points. Finally, the simplicity of the standard model enables a flexible and robust computational process, suitable for analysis of hundreds of species that vary enormously in their ubiquity, abundance, and population dynamics.

While the current approach produces trends that are useful for understanding population status of birds at regional or continental scales, the approach has a number of computational and spatial limitations. As implemented, it is a computationally intensive process, especially for wide-ranging species. This is due to the use of Markov chain Monte Carlo (MCMC) to estimate model parameters for relative abundance, and processing large MCMC chains to scale relative abundance to larger aggregate units. Additionally, their coarse resolution limits their ability to provide inference about local variation and processes. While trends can be scaled up to larger spatial units, they cannot be scaled down to smaller ones. The analytical stratum is the finest level of resolution, which limits the extent to which variation in trends can be attributed to processes occurring at finer spatial scales (Thogmartin et al. 2004, Bled et al. 2013).

Moreover, the current approach does not take full account or advantage of spatial relationships among counts. Modeling this structure would facilitate borrowing information across spatial boundaries, allowing more robust trend estimates in places where data are sparse (Waller and Gotway 2004, Blangiardo et al. 2013, Banerjee et al. 2014). Indeed, borrowing of information could possibly allow trends to be estimated at spatial scales that are finer than the spatial strata currently used (Thogmartin et al. 2004, Bled et al. 2013).

Previous work by Thogmartin et al. (2004), Bled et al. (2013), and Smith et al. (2015) offered spatially-explicit variations of the standard trend analysis approach for community science data. These works were focused on analysis of BBS data, but their approaches are easily related to analysis of CBC data. Instead of using the standard strata described above (Smith et al. 2015), Thogmartin et al. (2004) assigned count sites to irregular polygons, created by tessellation of BBS route locations. Bled et al. (2013) assigned routes to cells on a regular grid, with one-degree latitude and longitude spacing. All three studies utilized spatially-structured random intercepts for relative abundance per polygon, grid cell, or stratum. Thogmartin et al. (2004) utilized a fixed effect of year per polygon, but that effect did not incorporate spatial structure. Bled et al. (2013) and Smith et al. (2015) estimated relative abundances per year, and then trends were generated as derived parameters, as done in the standard analysis.

Here, we present a different approach for calculating temporal trends in relative abundance, one that takes advantage of the considerable spatial structure in CBC data. This approach borrows components from previous ones, incorporates new components that prioritize robust trend estimation at finer spatial scales, and employs a simplified and computationally efficient workflow. Similar to Bled et al. (2013), we assigned CBC count sites to cells on a uniform grid that covered North America. Like Thogmartin et al. (2004), temporal trends were explicit components of the spatial model. In contrast to previous work, effort and year effects were modeled as random slopes with spatial structure, following a spatially varying coefficient (SVC) approach (Gelfand et al. 2003, Finley 2011, Congdon 2014). Finally, unlike prior studies using MCMC, we used integrated nested Laplace approximation (INLA) to estimate Bayesian posteriors for model parameters (Lindgren and Rue 2015, Rue et al. 2017), which led to a dramatic decrease in computing time. The three goals of this report were to (*i*) describe an SVC approach to calculating trends in CBC data, (*ii*) employ the approach using data for the American Robin (*Turdus migratorius*), and (*iii*) compare trend results derived from the SVC approach to aggregate results derived from standard methods.

## Methods

### Statistical model

We modeled CBC counts for a given species, *y*_*i*, *k*, *t*_, for grid cell *i* encompassing count circle *k* during year *t*, as a random variable from a negative binomial distribution. Expected values for counts per grid cell, *μ*_*i*, *t*_ were assumed to be a function of spatially-structured grid-cell, count-effort, and year effects, plus unstructured variation among count circles. The linear predictor for *μ*_*i*, *t*_ took the form:

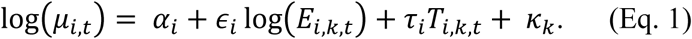

Parameters *α_i_* were modelled as cell-specific random intercepts with an intrinsic conditional autoregressive (CAR) structure (Besag et al. 1991). Given this structure, *α_i_* values came from a normal distribution, with a conditional mean related to the average of adjacent cells, and with conditional variance proportional to the variance across adjacent cells and inversely proportional to the number of adjacent cells. Spatial structure was incorporated into *α_i_* to allow for information about relative abundance to be shared across neighboring cells.

Parameters *ϵ_i_* were modeled as spatially-structured, cell-specific, random slope coefficients for the effort effect. These spatially varying coefficients (Gelfand et al. 2003, Banerjee et al. 2014, Congdon 2014) were also modelled with a CAR structure (Besag et al. 1991). Slopes were drawn from a normal distribution with a conditional mean related to the average of adjacent cells, and with conditional variance proportional to the variance across adjacent cells and inversely proportional to the number of adjacent cells. Spatial structure was incorporated into *ϵ_i_* to allow for information about the effort effect to be shared across neighboring cells. Effort was represented by *E*_*i*, *j*, *k*_, the number of party hours expended during a count, where a party hour was the count effort of one party of unspecified size for one hour. Pairing log-transformed counts with log-transformed effort in the linear predictor yielded a power function for effort correction, a flexible mathematical form that accommodated a decreasing, linear, or increasing impact of effort on expected counts (Butcher and McCulloch 1988, Link and Sauer 1999).

Parameters *τ_i_* were modeled as spatially-structured, cell-specific, random slope coefficients for the year effect. These spatially varying coefficients (Gelfand et al. 2003, Banerjee et al. 2014, Congdon 2014) were also modeled with CAR structure (Besag et al. 1991), where values came from a normal distribution, with conditional means and variances as described above. Spatial structure was incorporated into *τ_i_* to allow for information about the year effect to be shared across neighboring cells. Year, represented by *T*, was transformed before analysis such that max(*T*) = 0, and each preceding year took an increasingly-negative integer value. Given the scaling of effort and year variables, exp(*α_i_*) could be interpreted as a cell-specific expected count given one party hour of effort during the final year in the time series.

The final term in the model, *κ_κ_*, was an exchangeable random effect that accounted for variation in relative abundance among circles, possibly due to differences in habitat conditions or observer experience (Soykan et al. 2016). Note that the model did not include a normally-distributed, observation-level random effect to deal with overdispersed Poisson counts, i.e., *y*|*ε* ~ Poisson(*με*) and *ε* ~ normal(*μ*, *σ*), as is done for the standard approach (Sauer and Link 2011, Soykan et al. 2016). Rather, we used a negative binomial count distribution for *y*, i.e., *y*|*ε* ~ Poisson(*με*) and *ε* ~ gamma(*ϕ*^−1^, *ϕ*^−1^) (Linden and Mantyniemi 2011). These two approaches are expected to yield similar outcomes. However, as implemented in R-INLA, the latter approach returns a dispersion estimate while foregoing estimation of individual observation effects, which reduces computing time and the size of posterior samples.

### Case study

We developed a case study using data for the American Robin, from Audubon CBCs conducted across the continental US and Canada from 1966 through 2017, to demonstrate the SVC modeling approach and compare results with those using the standard approach. Before modeling the data, extreme outliers (> 3 SD from the mean, after log transformation) in counts and effort were removed. After filtering, there were 78,140 counts from 3,195 count circles over 52 years, for modeling.

Locations of the 3,195 unique count circles were mapped using the North American Albers Equal Area Conic projection (EPSG 102008, https://epsg.io/102008) and assigned to 880 cells on a grid divided along 100 km increments in latitude and longitude (Fig. 1A). Grid cells formed a continuous lattice within a non-convex polygon created using circle locations. A continuous uniform lattice was used to improve qualities of the neighborhood structure used in CAR modeling (Bled et al. 2013). The number of count circles per grid cell varied from 0 to 20, and averaged 2.43 (Fig. 1B). The number of neighbors for a given grid cell ranged from 1 to 8, and averaged 7.48. Note that grid cells with zero counts were retained during model estimation to preserve the spatial relationships between counts. However, before analyzing resulting trend estimates, cells with no observed counts were removed from the dataset, as we were not interested in interpolated trends for grid cells without CBC sites.

**Figure 1.**
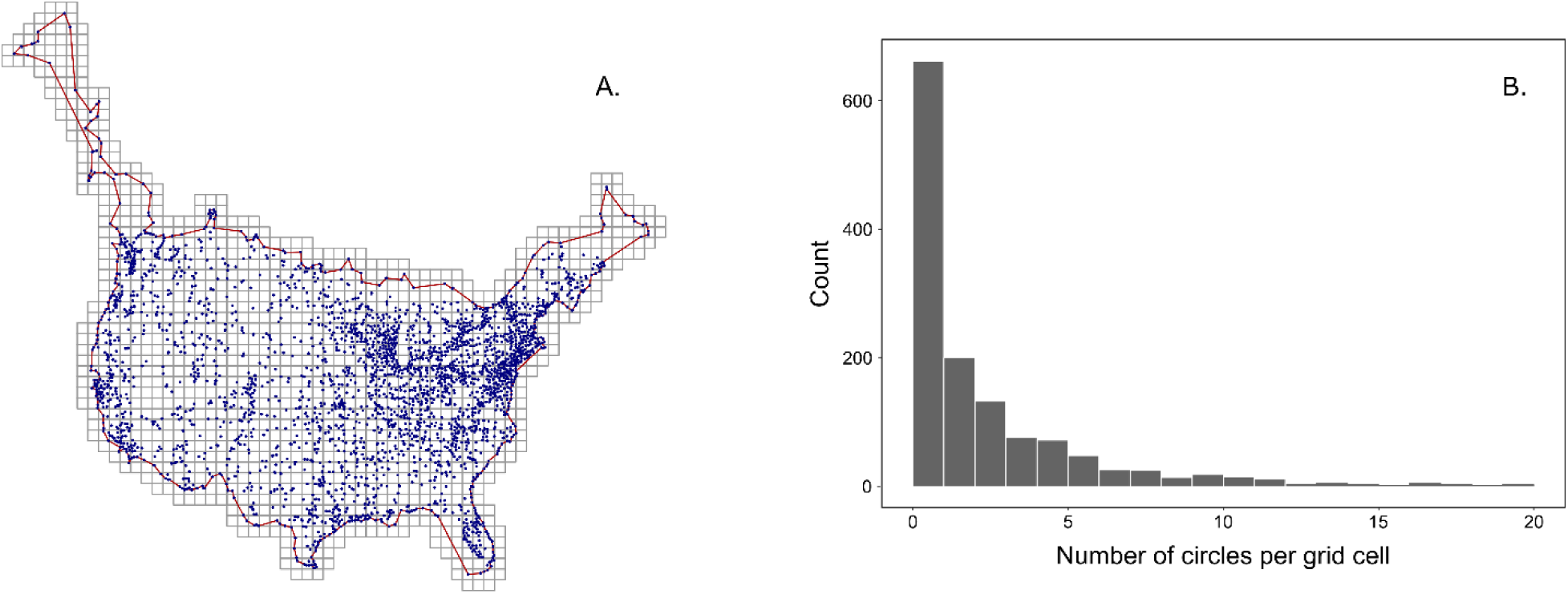
Grid cells used for spatial modeling of Christmas Bird Count data. (A) Cells were uniform, with 100 km sides, and were trimmed to a non-convex hull (red line) encompassing the count circle locations (blue circles). The number of count circles per grid cell (B) ranged from 0 to 20 and averaged 2.43. Cells with no circles were included during model analysis but removed for subsequent assessment of resulting trends (Figs. 2-4).

**Figure 2.**
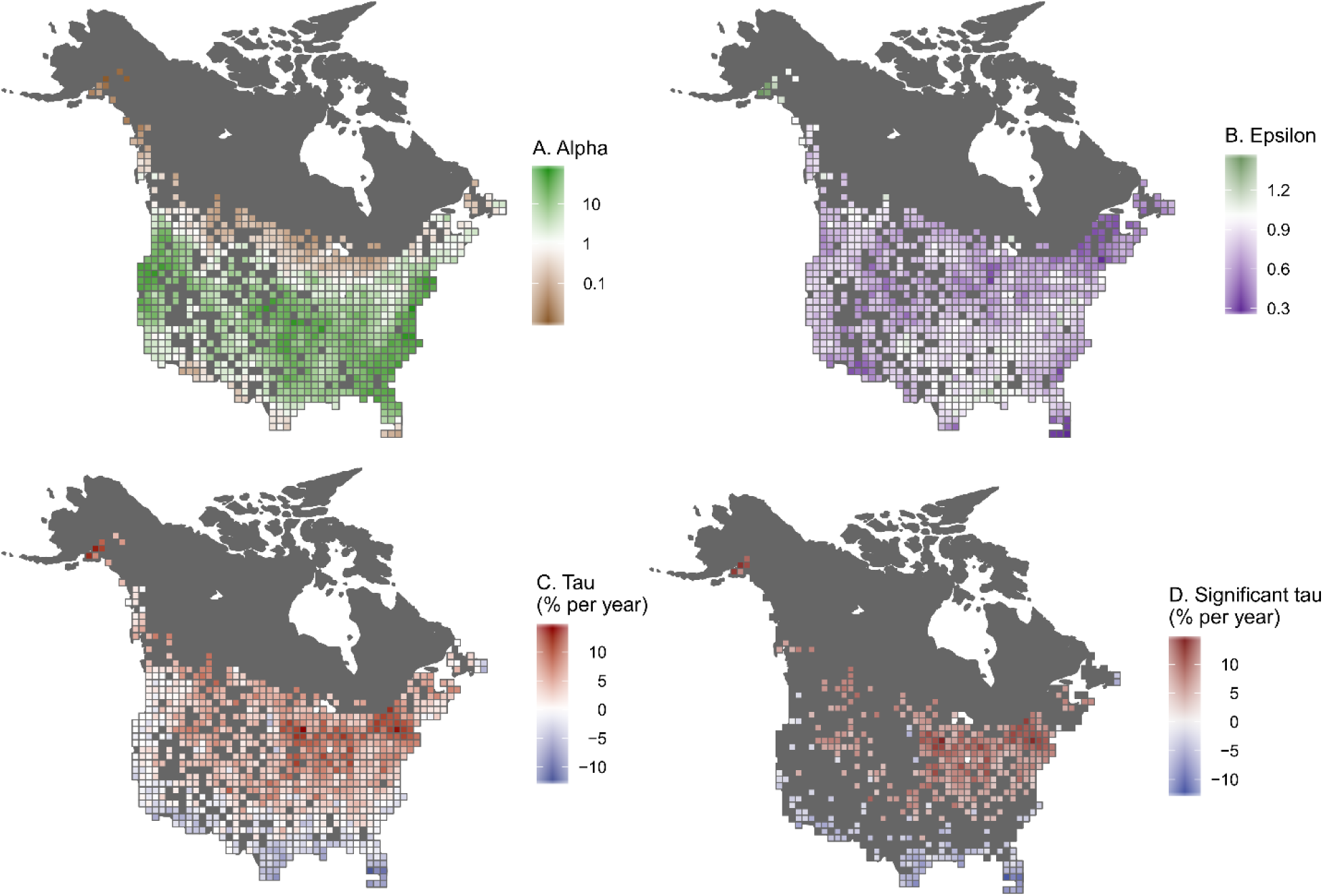
Maps showing spatial variation in posterior medians for parameters (A) *α_i_* (Alpha, relative abundance index), (B) *ϵ_i_* (Epsilon, effort-effect exponent) and *τ_i_* (Tau, long-term loglinear trend, percent change per year) per grid cell. For *τ_i_*, (C) all estimates and (D) only those significantly different from 0 are shown.

### Computing

The SVC model described above was analyzed in a Bayesian framework using the R-INLA package (Rue et al. 2017) for R statistical computing software (R Core Team 2016). Precision parameters for *α_i_*, *ϵ_i_*, and *τ_i_* random effects were assigned penalized complexity (PC) priors, with prior parameter values *U_pc_* = 1 and *a_pc_* = 0.01 (Simpson et al. 2017). Precision for the zero-centered, exchangeable, random circle effect, *κ*_κ_, was also assigned a PC prior with prior parameter values *U_pc_* = 1 and *a_pc_* = 0.01 (Simpson et al. 2017). The overdispersion term, *ϕ*, was assigned a PC prior with prior parameter value *l* = 7. Readers are referred to Simpson et al. (2017) for the details of, and rationale behind, PC priors, as well as the default structures and parameter values used in the R-INLA package.

Along with parameter estimates, R-INLA has the capacity to return from model analysis two values to evaluate individual model fit (Czado et al. 2009) and compare different models to one another (Gneiting and Raftery 2007, Link et al. 2017): cross-validation probability integral transform (PIT, Dawid 1984) and conditional predictive ordinate (CPO, Pettit 1990). For this application, we were not comparing multiple models. However, we extracted PIT values and visually inspected their histogram, as an approximate uniform distribution is expected for a model that fits the data reasonable well (Czado et al. 2009, Held et al. 2010).

Following model analysis, posterior medians and symmetric 95% credible intervals were computed per cell for *α_i_*, *ϵ_i_*, and *τ_i_*. Credible interval widths, representing estimate uncertainty, were computed by subtracting the lower credible limit from the upper credible limit per cell. Posterior summaries were then mapped to visualize spatial variation in 2017 abundance indices, effort effects, and 1966 through 2017 relative-abundance trends.

It is common, following CBC and BBS analyses, to aggregate trend information to larger scales that might be of interest to resource managers designing and implementing policies across states, provinces, BCRs, or nations (Sauer et al. 2003, Sauer and Link 2011, Soykan et al. 2016). After analysis of the SVC model for American Robin, we aggregated 100 km results to the BCR level in order to compare them to those produced using standard CBC analysis methods (Soykan et al. 2016). SVC trends were aggregated for each BCR by averaging trends for all equal-area grid cells where the cell centroid fell within the BCR. We evaluated the uncertainty around SVC trend estimates by comparing credible interval widths per cell to those calculated for a BCR using the standard approach. Aggregate trend estimates, along with their aggregate uncertainties, could also have be computed with R-INLA using functions for creating linear combinations. We did not employ these tools here because our main focus was on fine scale trends.

## Results

Model analysis using R-INLA took approximately 10 minutes to complete. Inspection of the PIT histogram indicate satisfactory model fit. The median of posterior medians for *a_t_* indicated that, on average, 4.28 robins were counted per party hour in 2017, but that number varied by several orders of magnitude across the species range, from 0.01 to 73.50. A map of posterior median values illustrated that the species was most abundant in regions the central part of their geographic range, and was least abundant along the northern and southern margins of their range (Fig. 2A).

Posterior median values for *ϵ*_i_, the power law exponent for the relationship between effort and counts, varied from 0.28 to 1.44, with a median value of 0.81. The 95% credible intervals for *ϵ_i_* indicated that 80% of estimates were not significantly different from 1, while all were significantly greater than 0. Estimates not significantly different from 1 indicated a positive linear relationship between effort and counts. Values significantly greater than 0 and less than 1 also indicated a positive relationship between effort and counts, but one with diminishing returns for additional count effort. A map of posterior median *ϵ_i_* values highlighted the spatial structure in the effort effect (Fig. 2B). Locations with posterior medians well below 1 were frequently locations with relatively low abundance indices (Fig. 2A), suggesting that the majority of robins in a count circle would be counted with relatively low effort. Locations with posterior medians closer to 1 were frequently locations with relatively high abundance indices (Fig. 2A), suggesting an endless supply of robins for CBC volunteers to count. The two parameters, *α_i_* and *ϵ_i_*, were significantly correlated across space, with a rank correlation coefficient of 0.26.

Posterior median values for *τ_i_*, the temporal trend from 1966 through 2017, when transformed to annual percent change, varied from −11.80 to 13.63, with a median value of 2.63. The 95% credible intervals for *τ_i_* indicated that 8% of estimates were significantly lower than 0, while 44% were significantly greater than 0. A map of posterior median *τ_i_* values (Fig. 2C) showed that trends in relative abundance had strong spatial structure. Credible intervals for *τ_i_* values were used to illustrate where trends were significantly negative or positive (Fig. 2D), showing that relative abundance during winter has generally decreased in the southern parts of their range and increased in the northern parts of their range. The parameters *α_i_* and *τ_i_* were significantly correlated across space, with a rank correlation coefficient of −0.16, indicating that the strongest trends were occurring at the margins of the geographic range where relative abundance was lowest.

The posterior median estimate for *ϕ*, the dispersion parameter, was exp[–log(0.55)] = 1.83, highlighting considerable overdispersion in American Robin counts relative to a Poisson distribution. Credible intervals for precision estimates for the random effects showed that all were important for explaining variation in the count data. When precision values were converted to a standard deviation scale, the random effects were ranked *α_i_* (SD = 1.58), *κ_κ_* (1.05), *ε_i_* (0.25), and *τ_i_* (0.04), in terms of the amount of variation explained in counts.

A common practice following standard CBC and BBS analysis is to aggregate trends from the analytical stratum level up to larger scales, such as the BCR level. Figure 3 shows the median of posterior median SVC trends across cells per BCR (Fig. 3A), along with the posterior median trend for each BCR from the standard analysis (Fig. 3B). Side-by-side visual comparison of these maps showed that aggregate trends were similar, regardless of method. The SVC approach gave a median trend of 2.11 across all BCRs, while the posterior median trend for the standard approach was 1.95 across all BCRs. Within BCRs, trend direction was consistent across the two methods in 28 of 32 BCRs. The rank correlation between BCR trends generated by the two methods was 0.88. Regarding differences, trends derived from the SVC approach changed more smoothly across the continent, as would be expected using a spatial statistical model. Also, the range of posterior median SVC trends (−4.58, 9.16) was slightly less than that for standard trends (−7.96, 14.26), especially near geographic range boundaries, as would be expected given the sharing of information across space.

**Figure 3.**
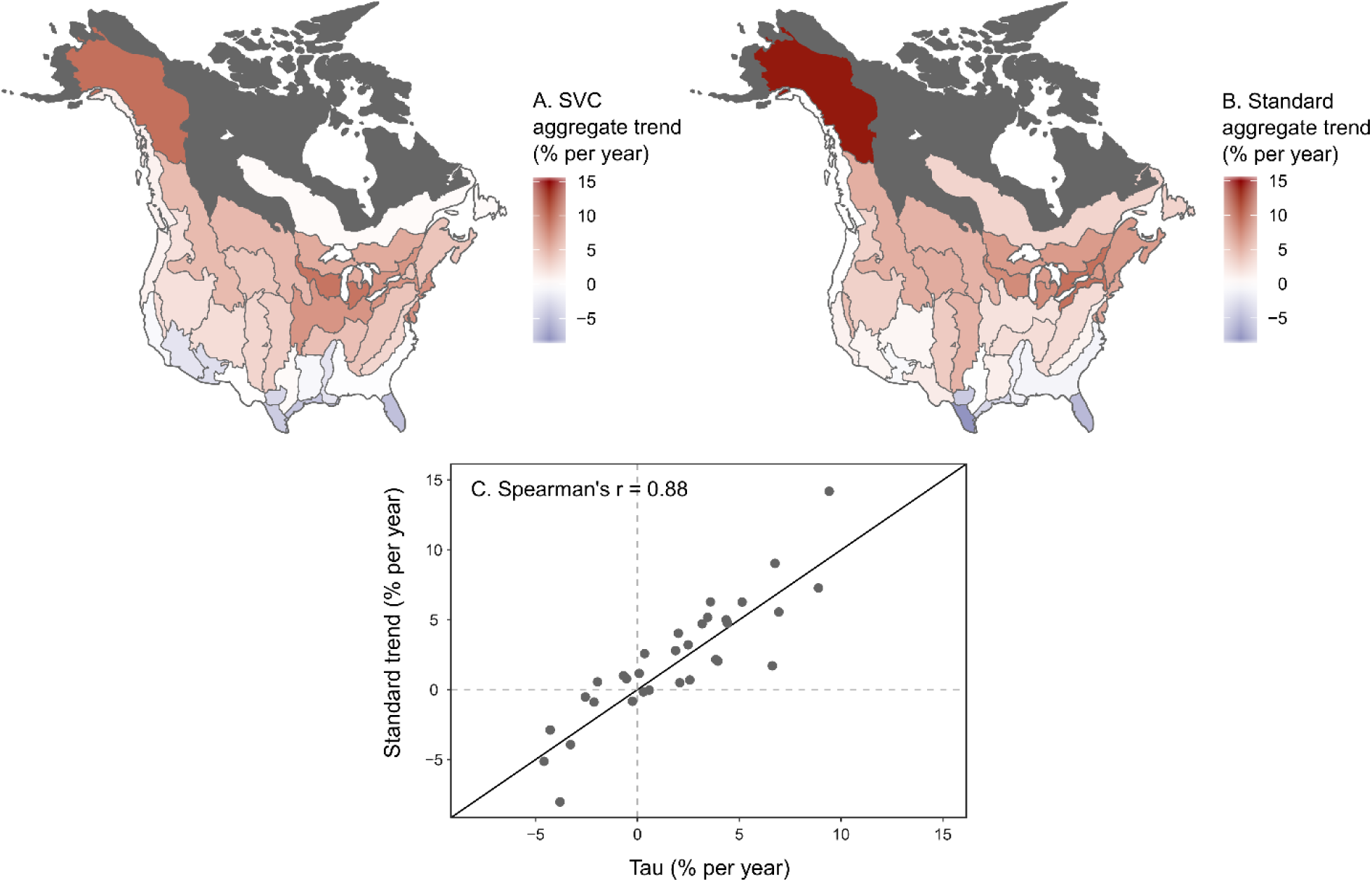
Comparison of posterior median trends, aggregated to Bird Conservation Regions, produced by (A) SVC and (B) standard methods, showing spatial variation in their relationships and their pairwise correlation (C). The dark grey diagonal line represents equality.

We also explored how the precision of trend estimates compared across the two approaches. Figure 4 compares the credible interval widths for SVC trends per grid cell (Fig. 4A) with those from the standard approach for aggregate BCR estimates (Fig. 4B). When compared to the standard approach, some SVC grid cells within a BCR, ones in information rich neighborhoods (Fig. 1A), had SVC trend estimates with remarkably narrow confidence intervals (Fig. 4C, SVC minimum). Other grid cells, ones in information poor neighborhoods (Fig. 1A), had trend estimates with relatively broad confidence intervals (Fig. 4C, SVC maximum). On average, however, interval widths of estimates per BCR were similar, regardless of method, if not slightly wider using the SVC approach (Fig. 4C, SVC median).

**Figure 4.**
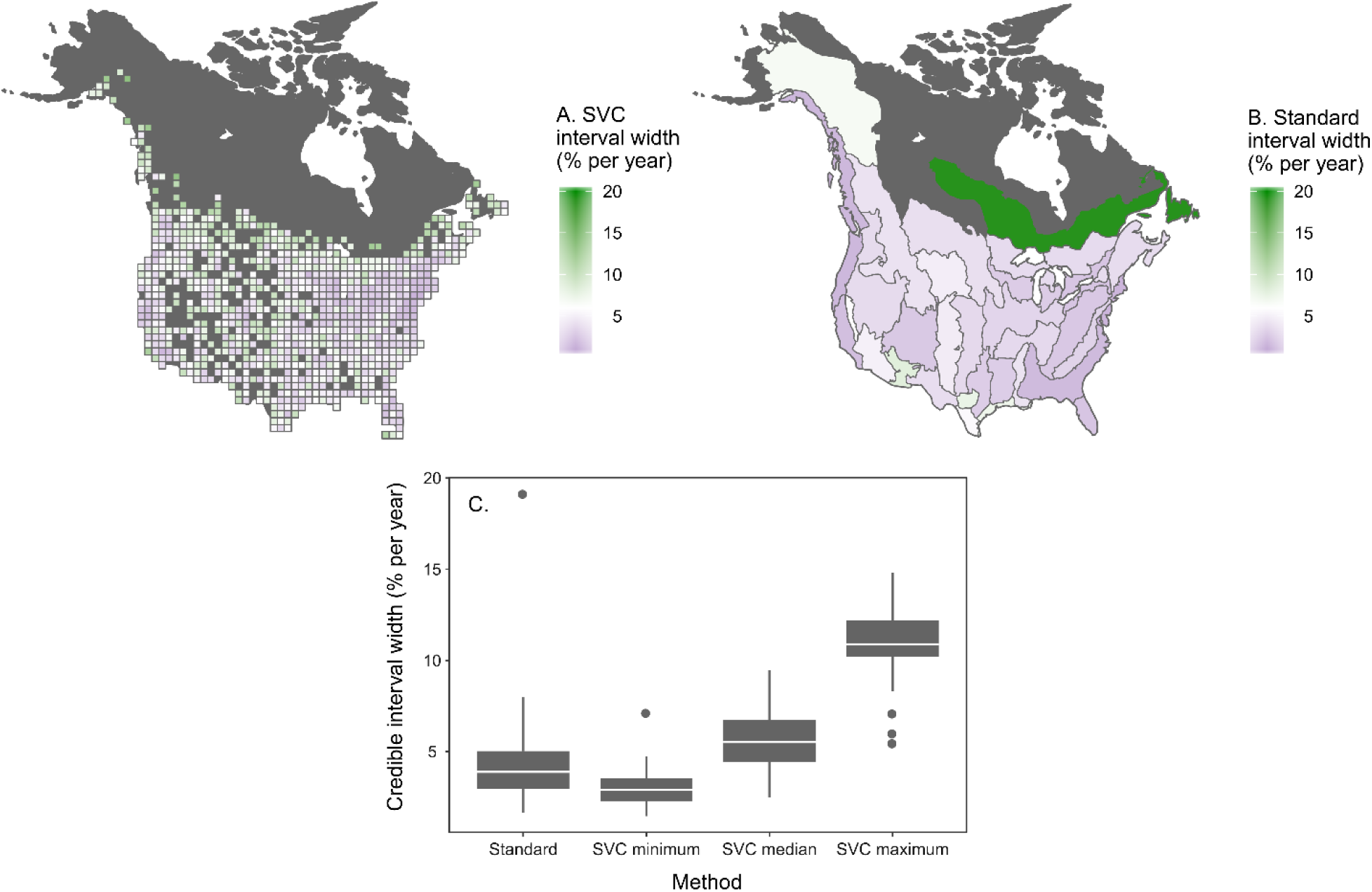
Comparison of 95% credible interval widths for (A) estimates of *τ_i_* (Tau, long-term log-linear trend, percent change per year per grid cell) from the SVC model and (B) analogous trends produced using the standard analysis and aggregated to Bird Conservation Regions, shown as maps and (C) summarized with box plots.

## Discussion

The goals of this analysis were to (*i*) describe a different approach for calculating trends from Audubon Christmas Bird Counts, (*ii*) demonstrate the approach using long-term count data for the American Robin, and (*iii*) qualitatively compare the trend results derived from the SVC approach to those derived using standard methods. We showed that the SVC approach generates trends at a finer spatial scale than the standard method, with comparable precision. Further, the SVC approach produced aggregate trends that were generally similar in direction, magnitude, and precision to those generated using standard methods.

To put resolution gains into context, consider that a CBC circle has a radius of approximately 12 km and an area of 452 km^2^. A 100 km grid cell, covering 10,000 km^2^, is approximate 22 times larger than a CBC circle. In comparison, the average analytical stratum has an area of 104,378 km^2^, approximately 231 times the area of a CBC circle. Thus, the SVC approach brought an order of magnitude increase in spatial resolution when compared to the standard approach. This increased resolution is expected to facilitate finer scaled investigations into the drivers of winter bird trends (Thogmartin et al. 2004, Bled et al. 2013, Smith et al. 2015).

Estimating trends at relatively high resolution was made possible by adopting spatial statistical techniques designed to borrow information across neighboring regions (Thogmartin et al. 2004, Bled et al. 2013, Smith et al. 2015). Employing spatial techniques also had implications for uncertainty in trend estimates. In the standard analysis, the uncertainty in a trend estimate depended upon the variation in trends across the circles within a stratum, and the number of circles in a stratum. In the SVC analysis, uncertainty depended upon those same two factors, but also depended upon those characteristics in the neighborhood of a grid cell. The consequences of this difference are demonstrated in Figure 4. In regions with many CBC circles (e.g., Piedmont BCR), SVC methods produced trend estimates with relatively low uncertainty (minimum credible interval width of 1.48) compared to the standard method (minimum credible interval width of 3.40), due to the density of information. Similar to Bled et al. (2013), we found that precision of SVC estimates also tended to be relatively high in regions at the edge of a species range where there were few counts (e.g., Boreal Softwood Shield BCR, maximum interval width of 12.02) when compared to the standard approach (maximum interval width of 19.14), due to borrowing of information across neighboring cells that crossed regional boundaries. In other parts of the continent with fewer, more isolated CBC circles (e.g., Southern Rockies Colorado Plateau BCR), the SVC methods produced trend estimates with relatively high uncertainty (minimum interval width of 3.47) compared to the standard method (minimum interval width of 2.80). It is not entirely clear if the small intervals of the standard approach are justified in this context. If the relatively few and far-between circles that fall within those large BCRs can be considered representative samples of that larger area, then estimates with high precision are reasonable, and certainly preferred. If it cannot be assumed that those circles are representative of the larger area, then estimating trends for smaller areas, in neighborhoods with more information, and basing uncertainty estimates on the amount of local information, seems more appropriate. Critical evaluation of this representative-sample assumption is particularly important when analyzing data from the CBC, because count site selection is not based on sampling design principles (Dunn et al. 2005), and count circles are neither randomly, nor evenly, distributed across the continent.

On a standard laptop computer, SVC model analysis using R-INLA took roughly 10 minutes for full Bayesian results. The standard approach, which employs MCMC, took approximately 10 hours for full Bayesian results on the same hardware. Had spatial statistical models been analyzed using MCMC, processing times would have been much longer. The difference in computing time was due to R-INLA producing highly accurate approximations of Bayesian posteriors, orders of magnitude faster than MCMC (Rue et al. 2009, 2017). The obvious benefit of shorter processing times is that, for a given set of computing resources, more time periods, more distinct model forms (e.g., Link and Sauer 2016), or more species can be evaluated. Even small differences in computing time add up when analyzing counts from tens of years, for hundreds of species, across thousands of count sites.

There were, as there usually are, tradeoffs for rapid model analysis. Specifically, R-INLA is an option for analysis whenever a statistical model can be expressed as a latent Gaussian model (Blangiardo et al. 2013, Rue et al. 2017). This was possible for the model used in this analysis. However, this would not have been possible had we chosen to use the effort-correction function developed by Link and Sauer (1999, 2006) and used in the standard analysis (Soykan et al. 2016). Here, we used a single-parameter, power-law function for effort correction because it could fit positive, negative, linear, increasing, and decreasing relationships (Butcher and McCulloch 1988) and was easily built into a latent Gaussian model. In contrast, the effort-correction function used for the standard approach is a two-parameter nonlinear function, which is more flexible and so will better-fit relationships that come to a rapid asymptote. Ideally, we would have tools for rapid analysis of spatial statistical models that incorporate the standard effort-correction function. In this choose-two situation, we erred towards rapid analysis of a spatial model with the simpler effort-correction function, because it allowed for more robust, if occasionally slightly biased (Link and Sauer 1999), estimates of the effort effect in regions where information was sparse. Robust estimates of effort effects are particularly critical when generating trends from CBC data, as count effort varies widely across time and space (Bock and Root 1981, Butcher et al. 1990, Dunn et al. 2005).

In addition to differences in model structure, there were other differences between the SVC approach outlined here and the standard approach. For instance, we chose to use a negative binomial count distribution, rather than a Poisson distribution with a normally-distributed, observation-level, random effect in the linear predictor (Link et al. 2006, Soykan et al. 2016). As described above, these two strategies should have similar outcomes, as a negative binomial distribution can be related to a Poisson distribution with a gamma-distributed, observation-level random effect added (Linden and Mantyniemi 2011). Given similar outcomes, we chose the negative binomial strategy as it resulted in many thousands fewer parameter estimates.

More generally, the SVC approach described here differed from the standard approach in that it was optimized, specifically, for computing long-term, log-linear trends in relative abundance at fine spatial scales. The emphasis on long-term, log-linear trends was motivated by requests from resource managers, who desire simple summary statistics that reflect overall population status for many species (Rosenberg et al. 2016, 2017). The emphasis on fine spatial resolution was motivated by requests from, both, researchers wishing to conduct research at relatively fine spatial scales, and Audubon Christmas Bird Count volunteers, who wish to learn how bird numbers have changed over the years in their local area. Given these two emphases, we did not incorporate additional model terms necessary for creating annual abundance indices. These indices are critical for those who wish to look beyond single, long-term trends, at detailed time series that give more information about the nature of abundance changes. Creating these annual indices is done by adding an additional random effect per cell and year, and combining these effects with *α* and *τ* Adding this effect to the SVC model is easily done in R-INLA. This effect could be specified as exchangeable, or have spatial or temporal structure. For this dataset, preliminary trials showed that adding an exchangeable effect to the model increased computing time to approximately 1 hour. We did not explore this model variant in depth because producing annual abundance indices was not a primary goal of this effort.

Despite the differences noted above, we learned that aggregate trends resulting from the SVC and standard approaches are similar in direction and magnitude. Precision at aggregate levels is generally similar, if not a bit lower with the SVC approach, due to different assumptions about how precision should, or should not, be related to the spatial distribution of counts. Our results suggests that an SVC approach can produce fine-scaled trends for some audiences, without paying a large price in precision, while producing aggregate trends for other audiences. These dual benefits, along with increased computational efficiency, make this SVC approach an attractive complement to the standard approach, one worthy of further exploration.

